# CorrAdjust unveils biologically relevant transcriptomic correlations by efficiently eliminating hidden confounders

**DOI:** 10.1101/2024.12.24.630258

**Authors:** Stepan Nersisyan, Phillipe Loher, Isidore Rigoutsos

**Affiliations:** Computational Medicine Center, Thomas Jefferson University, Philadelphia, PA 19107, USA

**Keywords:** CorrAdjust, correlations, confounders, PCA, mRNA, miRNA

## Abstract

Correcting for confounding variables is often overlooked when computing RNA-RNA correlations, even though it can profoundly affect results. We introduce CorrAdjust, a method for identifying and correcting such hidden confounders. CorrAdjust selects a subset of principal components to residualize from expression data by maximizing the enrichment of “reference pairs” among highly correlated RNA-RNA pairs. Unlike traditional machine learning metrics, this novel enrichment-based metric is specifically designed to evaluate correlation data and provides valuable RNA-level interpretability. CorrAdjust outperforms current state-of-the-art methods when evaluated on 25,063 human RNA-seq datasets from The Cancer Genome Atlas, the Genotype-Tissue Expression project, and the Geuvadis collection. In particular, CorrAdjust excels at integrating small RNA and mRNA sequencing data, significantly enhancing the enrichment of experimentally validated miRNA targets among negatively correlated miRNA-mRNA pairs. CorrAdjust, with accompanying documentation and tutorials, is available through GitHub at https://tju-cmc-org.github.io/CorrAdjust.

## INTRODUCTION

Confounder variables can lead to false positive and false negative calls in many types of statistical analyses of transcriptomic data. Often, such data contain known and latent variables that represent biological and technical factors and explain a significant fraction of gene expression variance [1–3]. Confounder correction is often overlooked when analyzing RNA-RNA correlations, a popular technique for the identification of co-expressed genes, gene prioritization, and even annotation of genes with poorly understood functions [4–6]. The methods for such correction are still in their infancy.

In differential gene expression analysis, confounders are routinely accounted for via their inclusion as covariates in generalized linear models such as DESeq2 [7] or edgeR [8]. The approaches that allow the estimation of hidden confounder factors in RNA-seq data include surrogate variable analysis (sva) [1] or removal of unwanted variation (RUV) [2]. In genome-wide association studies, principal components (PCs) are often used to adjust for the influence of ancestral heterogeneity when analyzing admixed populations [9, 10].

The existing methods for correcting transcriptomic correlation data can be broadly categorized into those that adjust for known variables and those that identify and remove latent factors. The most straightforward approach consists of residualizing known covariates from gene expression data with linear regression prior to computing the correlations, which is equivalent to computing partial correlations [11, 12]. A recently developed method named COBRA is also based on correcting for known variables: it decomposes the gene-gene covariance matrix into a linear combination of matrices, one per each confounder variable; the matrix associated with the intercept term represents the corrected covariances [13].

A common element of existing methods focused on identifying and removing hidden confounders is the estimation of latent variables through principal component analysis (PCA) or PCA modifications. In several such studies, “vanilla” PCs were directly residualized from the expression data [14–16]. Alternatively, PCA is applied only on a subset of genes with the lowest variance in an effort to maximize the likelihood of capturing unwanted variation instead of useful biological signals [17]. Yet another approach finds and removes hidden confounders after inferring a pattern from known covariates [18].

All these methods correct the expression data for a subset of “first N” PCs (or similar inferred variables). The choice of value for the parameter N critically influences the outcome. One approach removes all statistically significant PCs identified by the permutation procedure [14]. Other schemes estimate the number of components to eliminate by manually inspecting the distribution of correlations among positive controls (e.g., genes encoding ribosome subunits) and negative controls (e.g., a random set of genes) [15, 17], or by leveraging gene similarity matrices based on expression level correlations and Gene Ontology terms [16].

Previous studies used conventional machine learning metrics to evaluate pairwise correlations in the presence of reference sets of “positive” and “negative” gene pairs. These metrics include FDR (more widely known as “1 − precision”) [14], sensitivity, specificity, and area under the receiver operating characteristic curve (ROC AUC) [11, 12]. However, these metrics are not well-suited for evaluating gene-gene correlations due to typically low numbers of significantly correlated gene pairs, imperfection of reference gene sets, and difficulty in interpreting gene-level findings when evaluating all pairs together (see Methods).

CorrAdjust addresses these gaps by detecting and eliminating hidden confounders in correlation analysis. Key to the method is a new enrichment-based metric specifically tailored for evaluating correlations. In what follows, we present the new method, apply it to the computation of mRNA-mRNA and miRNA-mRNA expression correlations, and compare its performance to other schemes through extensive benchmarks using data from three popular public repositories.

## METHODS

### CorrAdjust: input data and problem formulation

In what follows, we will use the term “gene” to generically refer to transcripts, including protein-coding messenger RNAs (mRNAs) and microRNAs (miRNAs). Also, we will use the term “reference collections” to refer to *a priori*-defined, ground truth collections of gene sets. We will refer to gene pairs in which both genes belong to at least one common reference gene set as “shared genes.” Examples of reference collections include Gene Ontology (GO) [19] for mRNA-mRNA pairs (shared genes belong to at least one common GO term) and TarBase [20] for miRNA-mRNA pairs (shared genes are formed by miRNAs with their experimentally validated target mRNAs).

We consider a gene expression matrix *X* with *n* rows representing biological samples and *d* columns representing genes. We assume that the data is already normalized in a way that allows gene expression comparisons across samples. Optionally, the expression matrix could comprise several experimental techniques (e.g., mRNA-seq and small RNA-seq) with each gene labeled by its “gene type” (e.g., mRNA and miRNA). We focus on the array of Pearson’s correlation coefficients computed between expression levels of all possible gene pairs formed by input genes. In the case of multiple gene types, the analysis could be confined to correlations involving molecules of user-specified types only (e.g., only miRNA-mRNA correlations).

We rank the gene pairs in decreasing order of correlation, from most significant (highest rank) to least significant (lowest rank). Our objective is to determine whether the shared genes are overrepresented among the highly ranked pairs. Note that in some cases, it may make sense to assign the highest ranks to the gene pairs with the most negative correlations and *vice versa*. One such example is the case of miRNA-mRNA gene pairs, where interacting miRNAs and mRNAs are expected to be anti-correlated.

Two critical observations motivated us to develop a new metric.

<u>1. Only a small subset of highly ranked gene pairs is relevant.</u> Empirical evidence shows that most gene-gene correlations are centered around zero [4, 11, 12, 21, 22]. Thus, to

capture an enrichment signal, it is essential to consider only the subset of the significant, highly ranked correlations.

<u>2. Reference collections are not a *perfect* ground truth for co-expression studies.</u> By design, not all shared genes from reference collections are expected to have correlated abundances at the transcript level. For example, two genes can belong to the same GO term because they interact at the protein level, which does not imply any correlation between the corresponding mRNAs.

Considering these two points, we prioritize assessing whether highly ranked gene pairs tend to appear in the same reference sets (precision) over focusing on a fraction of shared genes that are correlated (recall, also known as sensitivity).

We define the highly ranked gene pairs by taking the “top *α*%” of the ranked pairs list. The value of *α* is explicitly set by the user. Doing so has the following key advantages. First, it does not depend on sample size, unlike thresholding which is based on correlation p-values. Second, correction for confounders often dramatically shrinks the distribution of correlations to zero (see Results), entangling the use of correlation threshold to define the set of significant gene pairs with uncontrolled changes in the set’s cardinality.

The parameter *α* controls the trade-off between precision and the number of gene pairs called significant. Indeed, it is closely related to the precision/recall trade-off since sensitivity is bounded from above by a multiple of *α* (Lemma 2 in Supp. Note S3). A small value of *α* would decrease the number of considered gene pairs and, thus, sensitivity, but potentially increase precision by selecting only the most significant correlations. On the other hand, a large value of *α* would select more pairs and increase sensitivity at the price of diluting precision by letting many uncorrelated pairs through. Note that the maximum precision achieved by a no-skill model equals the fraction of shared gene pairs out of all pairs, which is determined by the reference collection being used. Thus, maximizing precision is a challenge in itself. On the contrary, one can always achieve a sensitivity of 1 by setting *α* to 100%.

### CorrAdjust: metric for evaluating correlations in the presence of reference collections

Our approach comprises three steps (Figure 1). First, we rank gene pairs according to their correlation and mark the top *α*% entries. Second, for each gene, we compute overrepresentation scores by considering only gene pairs to which the gene belongs. Third, we average the overrepresentation scores from the previous step over all genes and all reference collections if more than one collection is available.

**Figure 1.**
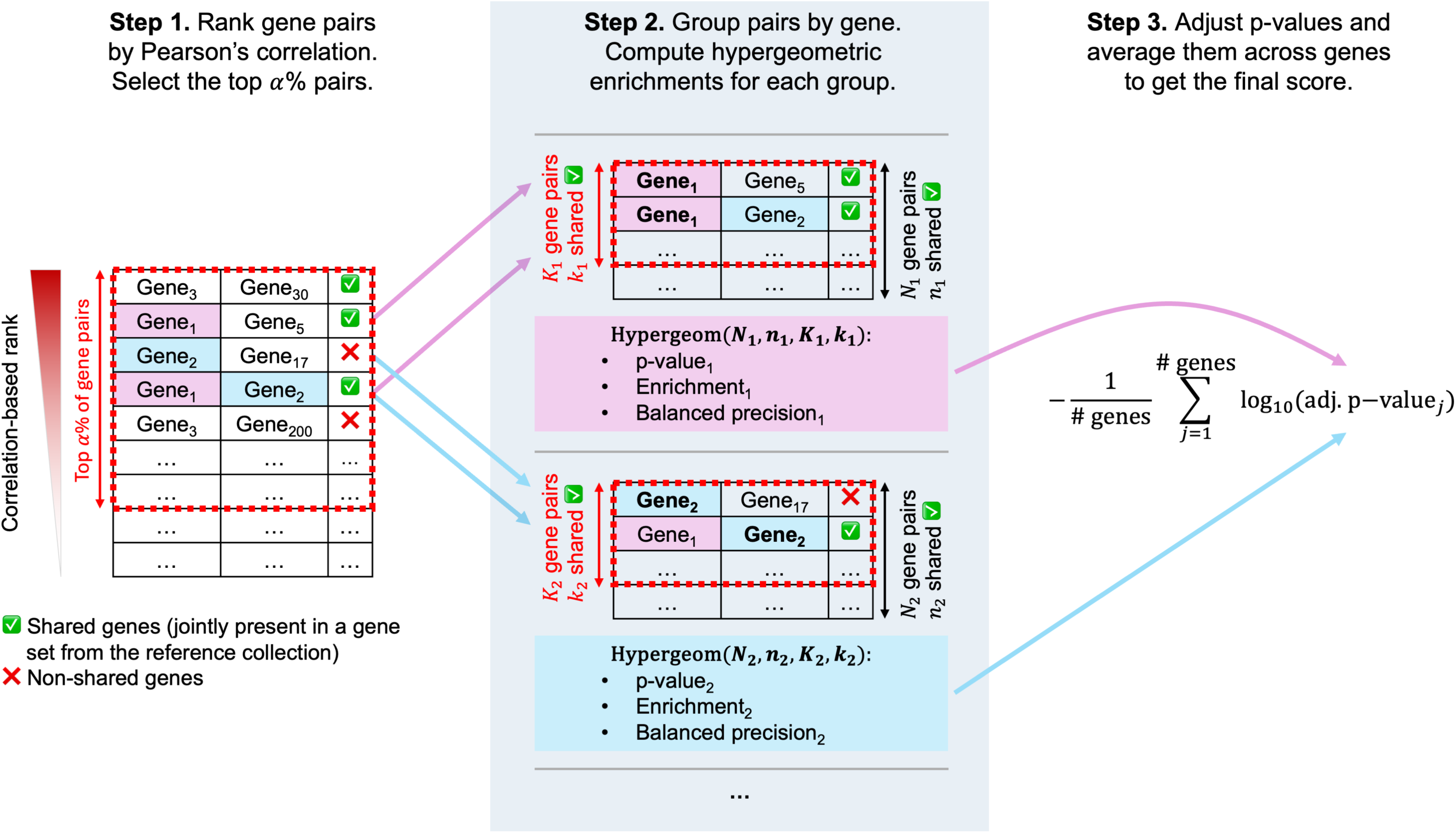
The outline of the novel metric for evaluating gene-gene correlations. In step 1, gene pairs are ranked according to their correlation value. Each gene pair is labeled as “shared” or “non-shared,” depending on whether the genes belong to at least one shared set from the reference collection. In step 2, gene pairs are grouped into blocks, each centered around one gene (pink and blue colors highlight example genes 1 and 2, respectively). Hypergeometric distribution-based statistics are then computed for highly ranked pairs in each block. Note that highly ranked gene pairs are defined based on the full table from step 1. In step 3, gene-wise p-values are adjusted for multiple testing, log-transformed, and averaged into a global enrichment score.

Let *K* denote the absolute number of ordered gene pairs associated with the *α*% cutoff, and *K*_*j*_ denote the number of pairs containing the *j*-th gene among the *K* pairs (*j* = 1, …, *d*). By definition, 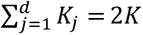, as the sum in the left part counts each gene pair twice. Let *k*_*j*_ denote the number of shared genes among the *K*_*j*_ pairs. By analogy, we define *N*_*j*_as the total number of gene pairs containing the *j*-th gene and *n*_*j*_ as the number of shared genes among the *N*_*j*_ pairs. In the simplest case of only one gene type, *N*_*j*_ = *d* − 1 for any *j*, as each gene form pairs with all other genes. In the case of two or more gene types, *N*_*j*_ values are not necessarily constant. For example, when miRNA-mRNA correlations are analyzed, *N*_*j*_ equals the total number of mRNAs or miRNAs if *j* represents miRNA or mRNA, respectively.

With *N*_*j*_, *n*_*j*_, *K*_*j*_, and *k*_*j*_ values computed, we assess the statistical significance of the overrepresentation of *k*_*j*_ shared genes among *K*_*j*_highly ranked gene pairs using the hypergeometric test. A commonly used statistic in this scenario is enrichment, enr_*j*_, which is defined as the ratio of *k*_*j*_ over its expected value under the null hypergeometric model *K*_*j*_ ∗ *n*_*j*_/*N*_*j*_. An undesirable property of enrichment is the dependence of its range of possible values on distribution parameters, thus making enrichment incomparable between the different genes (represented by index *j*). As a more suitable alternative, we adapted a recently developed balanced precision (BP) metric [23], which can be viewed as a non-linear monotonic transformation of enrichment (see Lemma 3 in Supp. Note S3):

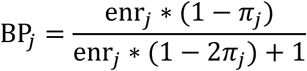

Here, *π*_*j*_ = *n*_*j*_/*N*_*j*_, a constant defined by the reference collection only. It is easy to see that BP_*j*_ = 0 corresponds to enr_*j*_ = 0, BP_*j*_ = 0.5 to enr_*j*_ = 1, and BP_*j*_ = 1 to the maximum possible value of enr_*j*_ = 1/*π*_*j*_. Thus, balanced precision could be directly compared between different genes.

The above computations become unstable when *K*_*j*_ is close to zero, i.e., when gene *j* participates in a small number of pairs from the top *α*% set. To tackle this problem, we use regularization inspired by the Bayesian beta-binomial model. Specifically, we add *v* = 5 “pseudo” gene pairs to *K*_*j*_ and *π*_*j*_ ∗ *v* gene pairs to *k*_*j*_ (the latter is rounded during hypergeometric p-value computation).

This procedure shrinks BP_*j*_ to 0.5 (or, equivalently, enr_*j*_ to 1) when *K*_*j*_is small, and the effect is negligible when *K*_*j*_ is large. We completely skip computations for genes with *N*_*j*_ < 1,000 to avoid problems with statistical power. For example, when integrating fewer than 1,000 miRNAs with mRNAs, we will not compute statistics for any *j* corresponding to an mRNA; computations will be carried out only for miRNAs (provided there are > 1,000 mRNAs in the dataset).

Lastly, we adjust the hypergeometric p-values across all genes using the Benjamini-Hochberg procedure. We define the global enrichment score by averaging − log_$%_(adj. p-value_*j*_) across all genes, similar to the approach used in Fisher’s combined probability test. The *j*-th p-value naturally combines *K*_*j*_ and enrichment into a single number, while neither can be used as a score alone. When several reference collections are available, we compute separate scores for each collection and then average the scores at the final step.

As an alternative to the enrichment-based metric, CorrAdjust offers the option to select “balanced precision at *K*” (denoted BP@*K*) – the BP computed over aggregated data 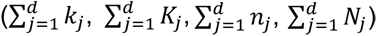. This can be viewed as a scaled version of “precision at *K*” (denoted P@*K*), a metric widely used in information retrieval within the machine learning community [24]. A recent study used a very similar quantity, false discovery rate (FDR), to evaluate mRNA-mRNA correlations (FDR = 1 − P@*K*) [14]. We strongly recommend using the global enrichment score as the default option, as it is significantly more informative and interpretable (see Results). BP@*K* and P@*K* are also vulnerable to Simpson’s paradox; specifically, comparing ratios of aggregated values 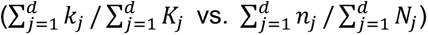 instead of comparing individual ratios per gene (*k*_*j*_/*K*_*j*_ vs. *n*_*j*_/*N*_*j*_) is a classical setup used to illustrate the paradox [25]. However, BP@*K* may prove valuable in scenarios not addressed in our study, such as cases where methylation and RNA-seq data are integrated and shared pairs represent promoters and their downstream genes, resulting in uniformly low *n*_*j*_values and an underpowered enrichment-based approach.

### CorrAdjust: PC correction procedure

The goal of our confounder correction procedure is to identify and regress out major hidden factors that drive undesired correlations between non-shared genes. Lemma 1 of Supp. Note S3 shows that PCs can be derived as orthogonal vectors that maximize the change in the covariance matrix’s Frobenius norm upon their residualization. This provides theoretical justification for leveraging PCs to correct the computed correlations.

We consider the transformation of *n* × *d* gene expression matrix *X* into *n* × *r* matrix of PCs, where *r* is the maximum number of PCs (*r* ≤ min (*n*, *d*)). We compute a partial correlation between expression levels of two genes *X*_*j*_, *X*_*l*_ controlling for a subset of PC_*j*_1__, …, PC_*j*_m__ by regressing-out PC_*j*_1__, …, PC_*j*_m__ separately from *X*_*j*_ and *X*_*l*_, and computing Pearson’s correlation on the residuals. The key observation is the mutual orthogonality of PCs, allowing to regress-out PC_*j*_1__, …, PC_*j*_m__ from columns of *X* consequentially in arbitrary order.

We use a greedy approach to find a set of PCs to regress out to maximize the global enrichment score defined in the previous subsection. We begin by limiting our search space to include only PCs that capture significant variation. In the current implementation, we use the permutation approach from the sva v3.5 package [1] (“num.sv” function). Other techniques can also be used, e.g., the “scree plot” method [26]. In the first iteration of the greedy algorithm, we loop over all PCs and select the one (PC_*j*_1__) with the maximum global enrichment score after its residualization.

In the *m*-th iteration, we loop over the remaining PCs and select the one (PC*_im_*) with the maximum global enrichment score after its residualization from expression data with PC*_i_*_1_ …, PC*_im-_*_1_ removed during the previous iterations.

### CorrAdjust: training and validation sets of <u>gene pairs</u>

Once a reference collection is selected, the identities of the shared genes emerge naturally. Despite PCA being independent of reference collections (unsupervised method), the procedure for selecting an optimal subset of PCs directly optimizes reference set enrichments and is supervised in nature. There are two elements that need to be evaluated to ensure generalizability and avoid overfitting: gene pairs and samples. CorrAdjust automatically splits *gene pairs* into 50% training and 50% validation sets. The reason there is no test set for gene pairs is the impossibility of creating one that is fully independent from the training set, as the same samples are used for PCA and correlation computations. Training and test sets of *samples* are described in the next subsection.

The training gene pairs provide the enrichment scores that are used as the objective function during the greedy optimization. In parallel, we evaluate the model using validation gene pairs at each greedy iteration. In a typical scenario, the optimization trajectory represents a quickly ascending curve that eventually reaches a plateau with much slower growth (see Results). We reach the early stopping point (i.e., the final number of PCs to remove) when both of the following criteria are met:

- The validation score improves by less than 5% over two consecutive iterations.
- The validation score differs from the global maximum score by less than 50%.

As the above early-stopping procedure uses two thresholds (5% and 50%), we recommend that users always visually assess whether the automatically determined peak is adequate (e.g., in rare cases, it may stop too soon) and manually refine it if needed.

### CorrAdjust: training and test sets of <u>samples</u>

We implemented an interface for using an optional test set of *samples* with CorrAdjust in the Python scikit-learn module style [27]. We use the training samples to derive gene-wise means for centering, PC loadings, and an optimal set of PCs. We then re-use these data to correct the test samples (if any) and evaluate their gene-gene correlations. We also implemented a training/test- friendly function for normalizing RNA-seq data using the median of ratios algorithm from DESeq2 [7] (vector of gene-wise geometric means is computed using the training set and reused with the test set).

### CorrAdjust: gene permutations for negative control runs

We also devised and made available an automatic permutation-based procedure to confirm the absence of overfitting. Specifically, we break the real link between gene expression and reference collections by randomly shuffling gene names of the expression matrix. At the same time, the vector of gene-gene correlations, PCs, and reference collections remain unchanged. This procedure can be used to demonstrate that CorrAdjust does not detect any statistically significant enrichments after correction.

### CorrAdjust: using the approach with samples from more than one condition

In some cases, input expression data comprise two or more groups (e.g., disease and control), and the user may be interested in performing differential correlation analysis [28, 29]. CorrAdjust includes built-in functionality to handle multiple user-defined sample groups. First, gene expression is centered separately in each group. Next, PCs are computed across all samples together. The standard CorrAdjust optimization procedure is then applied with a modified objective function. Specifically, correlations and global enrichment scores are computed for each sample group individually, and the scores are averaged across the groups. This approach ensures that the same PCs are residualized from each sample group, preventing the introduction of spurious differential correlations. At the same time, correlations are evaluated for each sample group independently, so CorrAdjust avoids removing real differential correlation signals.

### CorrAdjust: software implementation

We provide an implementation of CorrAdjust within a Python module available on GitHub (https://tju-cmc-org.github.io/CorrAdjust). Although this paper focuses on gene expression data, the CorrAdjust implementation uses the traditional machine terminology of “samples” and “features” in its interface, enabling users to apply it to any dataset.

We use the scikit-learn [27] implementation of PCA. The code is highly optimized and parallelized using Numba [30] and SciPy [31] modules and could easily handle datasets with hundreds of samples and 20,000 genes.

### RNA-seq data

We downloaded deep sequencing datasets from three different projects: datasets from The Cancer Genome Atlas (TCGA) generated from cancer patients, datasets from the Genotype-Tissue Expression (GTEx) project generated from healthy donors, and datasets from the Geuvadis consortium generated from lymphoblastoid cell lines (LCLs) of healthy participants to the 1000 Genomes Project. Supp. Table S1 shows summary of the data and the full lists of used samples and genes.

#### mRNA data

For TCGA and GTEx, we downloaded pre-mapped gene count tables from the GDC Data Portal (https://portal.gdc.cancer.gov; Data Release 32) and GTEx Portal (https://gtexportal.org/home/; GTEx Analysis V8), respectively. For Geuvadis, we downloaded raw sequencing reads in the FASTQ format (European Nucleotide Archive, PRJEB3366) and generated a read count matrix using STAR v2.7.10b [32] with GRCh38 reference genome and GENCODE v36 annotation. We selected only protein-coding genes for the downstream analysis.

#### Small RNA data

Small RNA-seq data are available only for the TCGA and Geuvadis samples. We downloaded raw FASTQ files for both projects and comprehensively profiled miRNA isoforms (isomiRs), tRNA-derived fragments (tRFs), rRNA-derived fragments (rRFs), Y RNA-derived fragments (yRFs), and their sequence variants using a slightly modified version of our recent pipeline [33]. Briefly, we used isoMiRmap [34] to map isomiRs, MINTmap [35] to map tRFs, and exhaustive exact mapping for rRFs and yRFs [33]. See Supp. Methods for a detailed description of the mapping pipeline.

#### Summary of included datasets

For TCGA, we used 8,981 primary tumor samples across 32 cancer types with both mRNA-seq and small RNA-seq data available (17,962 datasets). For GTEx, we used 6,295 samples across 11 healthy tissues. For Geuvadis, we used 403 samples with both mRNA-seq and small RNA-seq data available (806 datasets). We specifically excluded Geuvadis datasets generated by laboratory “number six” as they used more sequencing cycles in small RNA-seq than the rest of the datasets. We also excluded datasets that corresponded to technical replicates.

#### Data normalization

We separately processed each TCGA cancer type, GTEx tissue type, and Geuvadis data. We randomly split samples into 50% training and 50% test sets. We kept only mRNAs with a median abundance ≥ 1 transcript-per-million (TPM) and small RNAs with a median abundance ≥ 10 reads-per-million (RPM). We normalized filtered raw read count matrices with the median of ratios algorithm [7], separately for mRNA-seq and small RNA-seq. Then, we combined mRNA and small RNA matrices (TCGA, Geuvadis) and winsorized abundances of each feature (mRNA or small RNA) using lower 5% and upper 5% expression values. Finally, we applied the log_(_(*x* + 1) transform to the winsorized and normalized counts. We used only training samples to derive all the parameters used for filtering (median TPM or RPM), median of ratios normalization (gene-wise geometric means), and winsorization (expression quantiles).

### Reference collections

#### mRNA-mRNA pairs

We downloaded reference gene sets in the GMT format from MSigDB v2023.2 [36]. We used only Canonical Pathways and Gene Ontology (GO) as reference collections for running CorrAdjust.

#### miRNA-mRNA pairs

We used two reference sets for miRNA-mRNA pairs: experimentally validated miRNA targets from TarBase v9.0 [20] and purely sequence-based predicted miRNA targets by RNA22 v2.0 [37]. In both cases, we used only a subset of canonical isomiRs [38].

### Alternative confounder correction methods

We compared CorrAdjust with several other methods that also correct for hidden and known confounders.

#### sva_network

Parsana et al. [14] suggested residualizing the “first N” statistically significant PCs from gene expression data before computing correlations. At the time of running these benchmarks (October 2024), the publicly available implementation of this approach (“sva_network” function from the sva package) incorrectly centered the input data before running PCA. Because of this, we only used the sva package to estimate the number of PCs to regress out (“num.sv” function) and then used CorrAdjust for the actual residualization, mimicking sva_network behavior. We ran a PCA using the training samples and evaluated the corrected correlations using the test samples.

#### Linear regression on known confounders

We used multiple linear regression to residualize known variables from expression data. The variables included:

- sex, race, and age for the TCGA datasets;
- sex, age, RNA integrity number (RIN), and post-mortem interval (PMI) for the GTEx datasets;
- sex, population, and laboratory number for the Geuvadis datasets.

We fit regression models using training samples only. We discarded samples with missing information (TCGA only). We recomputed CorrAdjust scores on the remaining samples (while keeping PCs the same) to ensure comparability.

#### COBRA

We ran a Python implementation of the COBRA tool [13] from the netZooPy v0.10.6 package using the same known confounder variables. Since the tool does not provide an intuitive way of using training and test samples, we directly fit COBRA on the test samples.

## RESULTS

CorrAdjust is a novel method for identifying and correcting hidden confounder variables in transcriptomic correlation studies. It uncovers hidden confounders among the PCs of the gene expression matrix based on our theoretical proof that PCs are the factors with the greatest impact on the gene-gene covariance matrix upon their residualization (Lemma 1 in Supp. Note S3). CorrAdjust identifies a subset of confounder PCs by maximizing the enrichment of gene pairs that are jointly present in one or more gene sets from the reference collection among the top *α*% of the most correlated pairs. For simplicity of notation, we will use the term “shared genes” to refer to gene pairs that are jointly contained in the reference collection.

Key to CorrAdjust is our novel metric for evaluating enrichment of shared genes among the highly correlated pairs. Figure 1 outlines the steps constituting the computation of our new metric. First, the gene-gene pairs are ranked according to the correlation of their expression values. The top *α*% entries of the ranked gene pair list are then marked as “highly ranked”. Next, the array of all gene pairs is segmented into blocks, with the *j*-th block corresponding to all gene pairs formed by the gene with index *j*. Within each block, we use the hypergeometric distribution framework to quantify whether shared gene pairs are overrepresented among the highly ranked ones. Finally, p-values derived for each gene are adjusted for multiple testing, and negative logarithm values of adjusted p-values are averaged to generate a global enrichment score.

In the next two subsections, we will apply CorrAdjust to real RNA-seq data.

### Case 1: correcting mRNA-mRNA correlations in whole blood datasets from GTEx

After filtering for low-expressed genes, the data contain expression levels of 10,021 protein-coding genes across 755 samples. We further split the samples into training (n=378) and test (n=377) sets and 50,205,210 gene pairs into equal training and validation sets. There are 19 statistically significant PCs according to the permutation test (see Methods).

We ran CorrAdjust using two reference collections: Canonical Pathways and Gene Ontology (GO). Figure 2A shows optimization curves for the training set of samples, with the Y-axis showing the global enrichment score for Canonical Pathways, GO, and their average. The numbers on top of the curves show the identities of PCs selected at each iteration. After 3 rounds of greedy selection, the score reaches a plateau, and PC1, PC2, and PC3 are selected for elimination. Canonical Pathways and GO scores follow a very similar pattern, suggesting that both collections capture a common enrichment signal. Importantly, scores computed using training and validation *gene pairs* are almost identical, showing a complete absence of overfitting with respect to gene pairs. Next, we evaluated the same trajectory of PC correction using the test set of *samples* (all gene pairs together). The test scores closely resemble training scores (Supp. Figure S1A).

**Figure 2.**
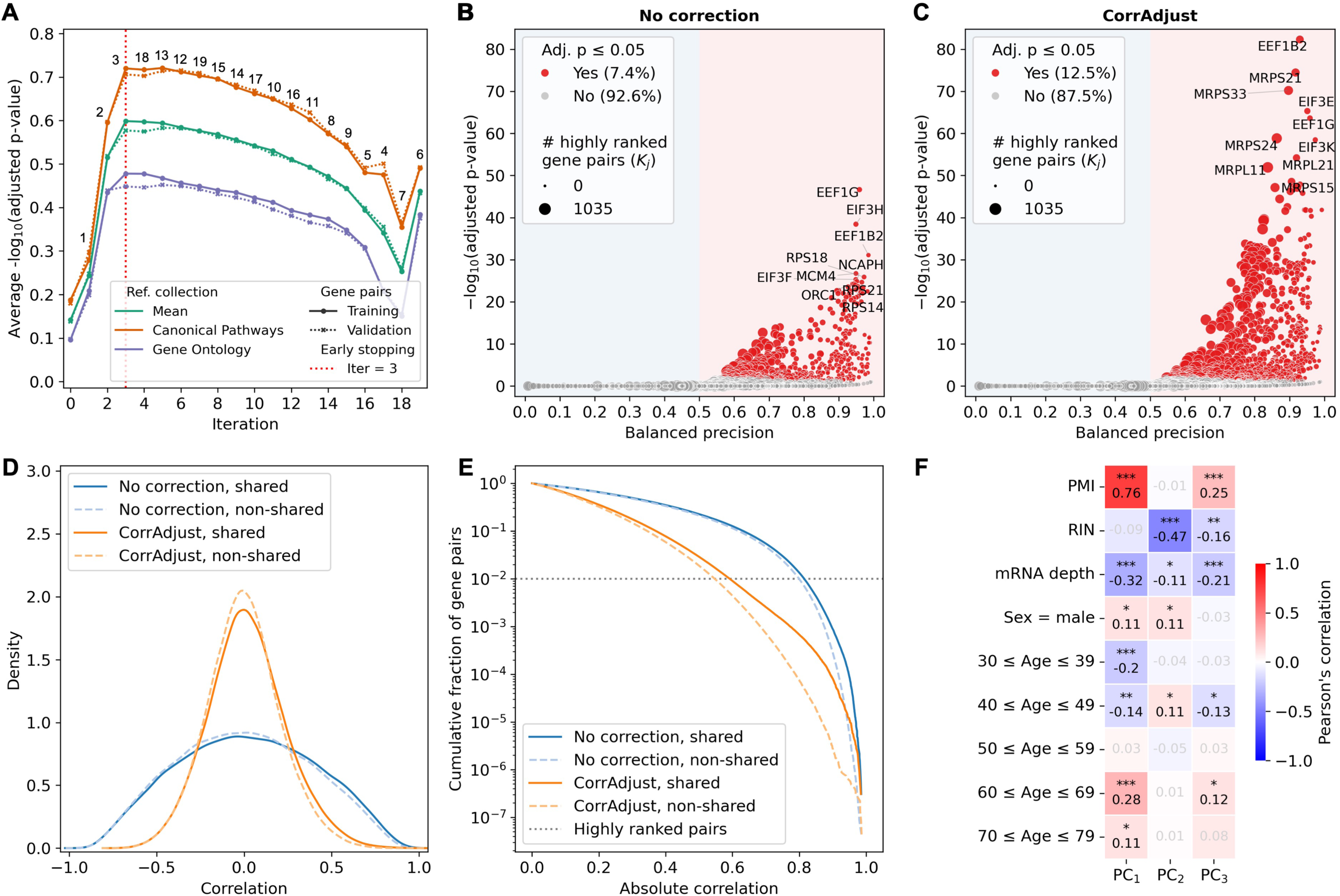
Application of CorrAdjust to mRNA-mRNA correlations computed over GTEx whole blood dataset. (A) Optimization trajectory of CorrAdjust (training samples). Each iteration (X-axis) corresponds to the selection of one PC (number on top of the curves). Iteration 0 corresponds to uncorrected data. The Y-axis shows global enrichment scores for Canonical Pathways, Gene Ontology, and the average of these scores. (B-C) Volcano plots before and after CorrAdjust correction (test samples, all gene pairs, Canonical Pathways). Each marker represents one gene. The marker area stands for the number of highly ranked gene pairs involving the corresponding gene (top *α*% approach, see Methods). The top 10 genes by adjusted p-value are annotated. (D-E) Kernel density estimation and cumulative distribution function of gene-gene correlations before and after CorrAdjust correction. Solid lines correspond to the pairs of mRNAs that are shared by at least one Canonical Pathway. (F) Correlations between identified confounder PCs and known covariates (training samples): post-mortem interval (PMI), RNA integrity number (RIN), sequencing depth, sex, and age group.

Figure 2B and Figure 2C show gene-level Canonical Pathways enrichment scores before and after correction, respectively (test set of samples, all gene pairs). Each dot represents an individual gene and depicts its three main quantities:

- the Y-axis represents the adjusted p-value (a dot is colored in red if the adjusted p-value is ≤ 0.05);
- the marker size represents the number of significantly correlated genes with the corresponding gene (*K*_*j*_ in the notation of Figure 1 and Methods);
- the X-axis represents balanced precision.

As it is evident from the plots, the removal of PCs strongly increases pathway enrichments across many genes.

Figure 2D shows the global impact of PC correction: gene-gene correlations are significantly shrunk toward zero. Together with increased enrichment scores, such shrinkage indicates the removal of a significant source of unwanted variance, driving high false positive correlations. Correlations of gene pairs shared by at least one Canonical Pathway (solid lines) are, on average, higher than for non-shared gene pairs (dashed lines). Figure 2E provides more insight into the distribution’s tails on a cumulative logarithmic scale. After correction, the difference between shared and non-shared gene pairs becomes noticeably more pronounced (note how the gap between solid and dashed lines increases toward higher correlations). The dotted horizontal gray line shows the used *α* = 1% threshold; before correction, it corresponds to the absolute correlation of 0.8 and becomes 0.55 after correction.

Some of the residualized PCs could be clearly interpreted (Figure 2F and Supp. Figure S2). For example, PC1 differentiates between samples obtained post-mortem (positive PMI) and pre-mortem (negative PMI), which was previously reported for the same GTEx whole blood dataset [39]. PC2 and PC3 also have statistically significant associations with known variables (PMI, RIN, sex, age), albeit with much weaker effect sizes. Below, we will show that adjusting for known confounders results in worse scores than CorrAdjust.

### Case 2: correcting miRNA-mRNA correlations in LCL datasets from Geuvadis

For the second example, we ran CorrAdjust on 403 LCL samples for which both mRNA-seq and small RNA-seq datasets are available. This time, we used the following reference collections: the miRNA-mRNA pairs of TarBase [20] (experimentally validated miRNA targets) and the miRNA-mRNA pairs generated by RNA22 [37] (sequence-based computationally predicted miRNA targets). Since miRNAs downregulate their target genes, we assigned the highest ranks to the most negative miRNA-mRNA correlations.

CorrAdjust selects 8 PCs to be eliminated from the data: PC1, PC3-PC6, PC9, PC10, and PC12 (Figure 3A). There is no evidence of overfitting both from the validation pairs (Figure 3A) and the test samples (Supp. Figure S1B) perspectives. Strikingly, in the absence of any correlation correction, none of the miRNAs show a statistically significant overrepresentation of predicted targets among anti-correlated miRNA-mRNA pairs, for either the TarBase or the RNA22 reference sets. However, after correlation correction, most miRNAs show statistically significant enrichment: 78% of miRNAs for TarBase and 60% of miRNAs for RNA22. Figures 3B-C show volcano plots with TarBase miRNA target enrichments before and after the CorrAdjust application.

**Figure 3.**
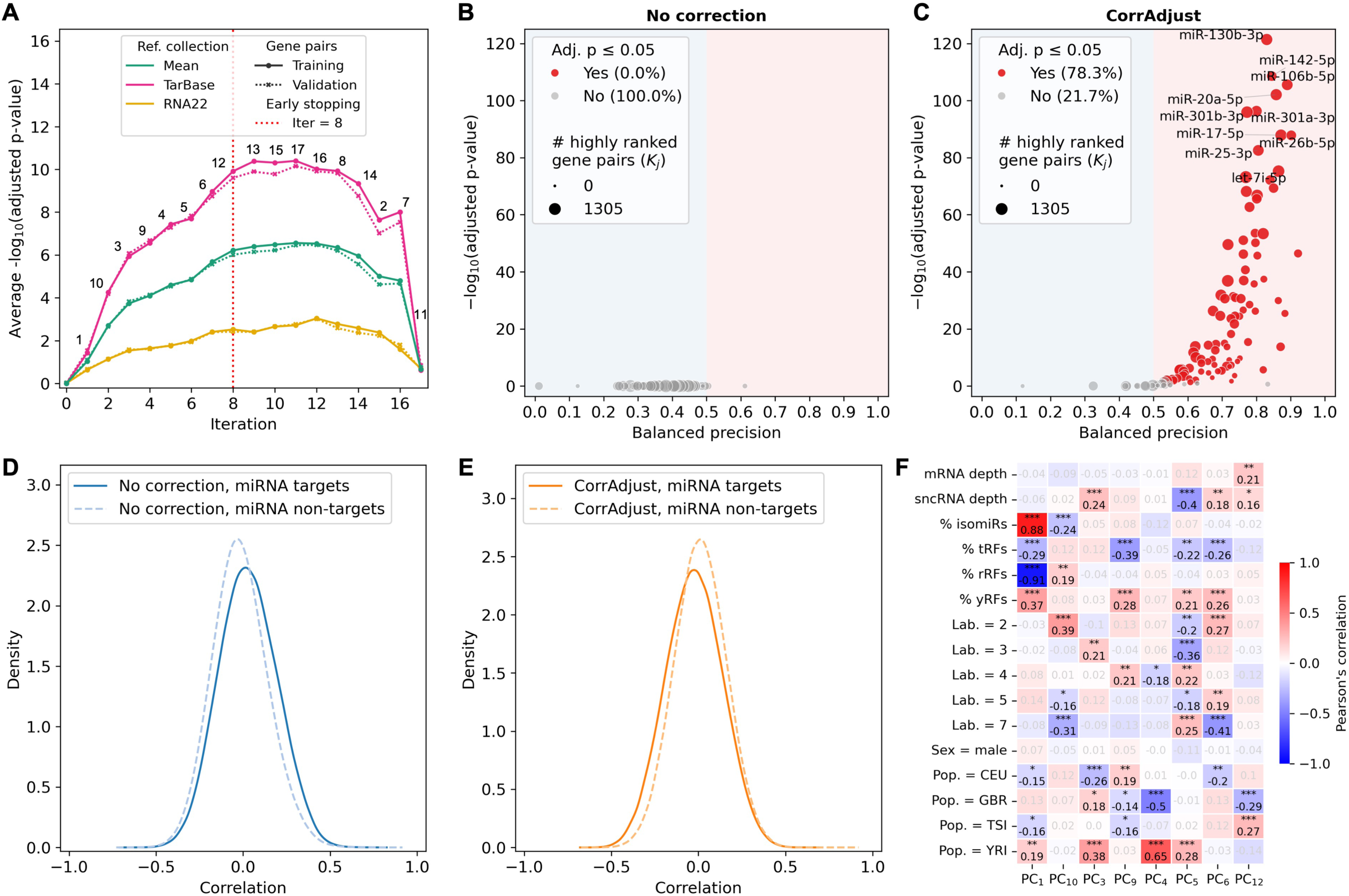
Application of CorrAdjust to miRNA-mRNA correlations computed over Geuvadis LCLs dataset. (A) Optimization trajectory of CorrAdjust (training samples). Each iteration (X-axis) corresponds to the selection of one PC (the number on top of the curves). Iteration 0 corresponds to uncorrected data. The Y-axis shows global enrichment scores for TarBase, RNA22, and the average of these scores. (B-C) Volcano plots before and after CorrAdjust correction (test samples, all pairs, TarBase). Each marker represents one miRNA. The marker area stands for the number of highly ranked miRNA-mRNA pairs involving the corresponding miRNA (top *α*% approach, see Methods). The top 10 miRNAs by adjusted p-value are annotated. (D-E) Kernel density estimation of miRNA-mRNA correlations before and after CorrAdjust correction. Solid lines correspond to pairs of miRNAs and their experimentally validated targets from TarBase. (F) Correlations between identified confounder PCs and known covariates (training samples): mRNA sequencing depth, small non-coding RNA (sncRNA) sequencing depth, composition of sncRNA-omes, laboratory ID, sex, and population.

This dramatic improvement could also be noted at the level of whole miRNA-mRNA correlation distributions (Figure 3D-E). Before the CorrAdjust correction, correlations between miRNAs and their experimentally validated targets (solid blue line) tend to be more positive than for non-targets (dotted blue line), contradicting the biological knowledge. The correction flips this pattern, putting experimentally validated pairs (solid orange line) to the left of non-targets (dotted orange line). Supp. Figure S3 shows intermediate states between uncorrected and fully corrected data. Adjusting for PC1 removes the positive gap between the correlation distributions of miRNAs with their targets and non-targets, while adjusting for the subsequent PCs gradually creates the expected negative gap. Interestingly, PC1 essentially represents % of isomiRs or % of rRFs in the small RNA-seq datasets (Figure 3F), suggesting that the positive bias of miRNA-mRNA correlations in Figure 3D is primarily driven by unwanted variance in the molecular composition of the small RNA datasets. The subsequent PCs have weak to moderate associations with both technical variables (sequencing depth, laboratory) and donor populations (Figure 3F).

### CorrAdjust outperforms other methods when correcting miRNA-mRNA correlations

We evaluated the ability of CorrAdjust and several competing methods to improve enrichments of TarBase and RNA22 targets among negative miRNA-mRNA correlations using 8,981 primary tumor samples spanning 32 TCGA cancer types. The competing methods included the sva_network method [14], correction for known factors with linear regression (see Methods for the list of factors), and correction for known factors with COBRA [13]. As a negative control experiment, we ran CorrAdjust on expression tables with shuffled mRNA names. We used *α* = 5% threshold for all datasets (Supp. Note S1 describes the selection of *α*).

Figure 4 shows the benchmarking results computed over test samples and all miRNA-mRNA pairs: the left panel compares CorrAdjust with hidden confounder correction methods, and the right panel compares it with known factor correction approaches. In virtually all cases, CorrAdjust outperforms the other techniques and shows a striking increase in global enrichment score (X-axis) compared to uncorrected correlations.

**Figure 4.**
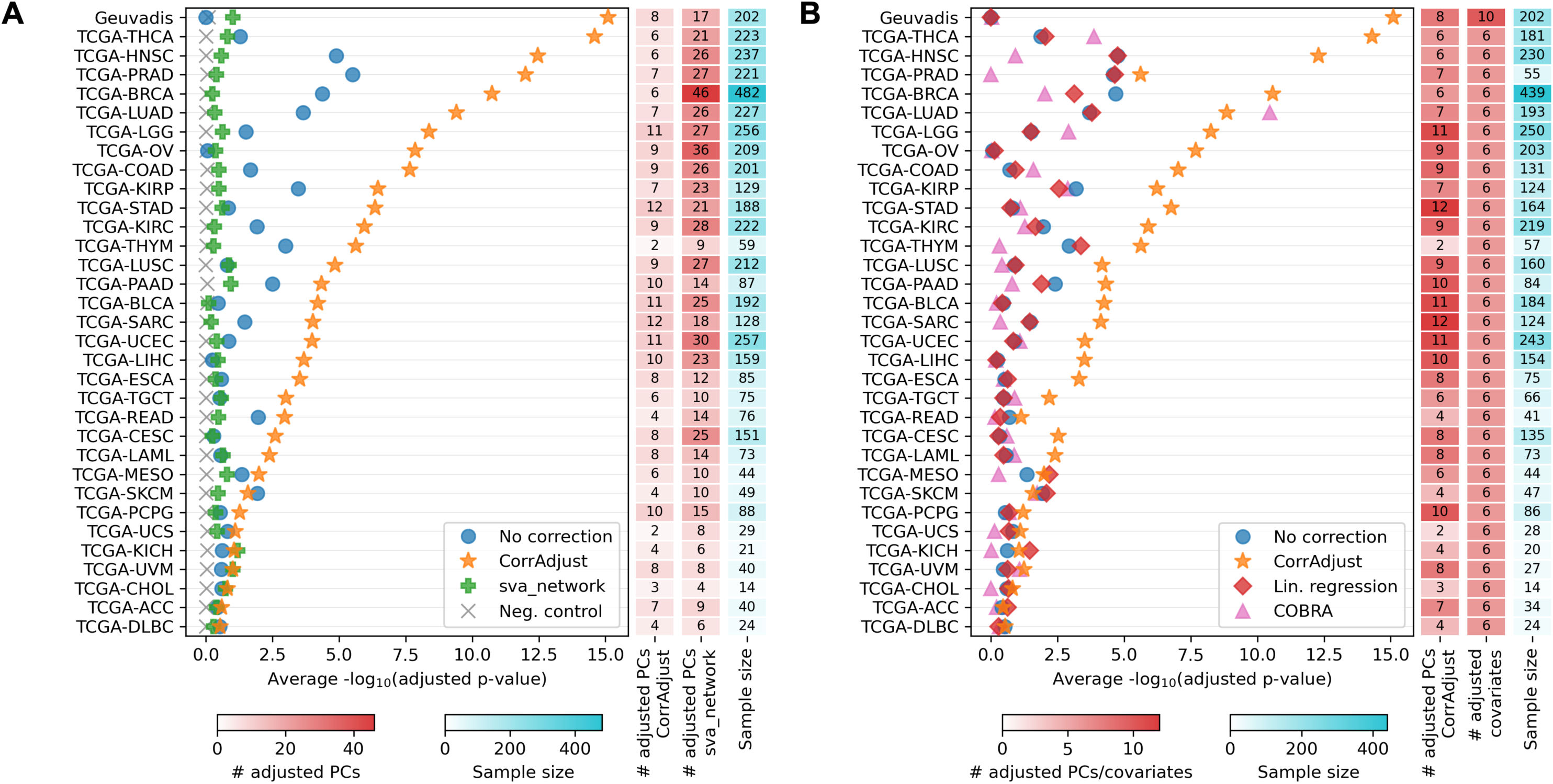
Benchmarking results of CorrAdjust and alternative methods using miRNA-mRNA correlations (test samples, all pairs). (A) Methods for hidden confounders correction. (B) Methods for known covariates correction. The X-axis shows the global enrichment score (average of TarBase and RNA22). The columns to the right of the main plots show the number of PCs adjusted by CorrAdjust, the number of PCs adjusted by the sva_network approach, the number of adjusted known covariates, and the number of samples in the training set.

The median number of PCs regressed out by CorrAdjust is 8. Even though the sva_network approach removes significantly more PCs (median = 18), it results in lower than raw scores in 24/33 datasets (Figure 4A). Known factor correction approaches also perform poorly, with linear regression and COBRA generating *worse* scores in the 16/33 and 21/33 datasets, respectively (Figure 4B). The same trends persist with scores computed over training samples and training miRNA-mRNA pairs (Supp. Figure S4A-B), showing that the low performance of these methods is not due to poor generalizability from training to test samples. Interestingly, the sva_network “first N” approach shows non-stable behavior with respect to the number of PCs removed, so it cannot be improved by simply regressing out fewer (but predefined) number of PCs (see representative examples in Supp. Figure S4C-E).

Importantly, CorrAdjust training and test sample scores are extremely correlated (Spearman’s *r* = 0.96, *p* = 2.25 ∗ 10^−19^), as well as the training and validation miRNA-mRNA pair scores (*r* = 0.95, *p* = 1.17 ∗ 10^−16^). Additionally, all negative control experiments show near-zero scores even when evaluated with training samples and training gene pairs (Supp. Figure S4A). Taken together, there are no signs of overfitting when conducting supervised PC selection with CorrAdjust.

There is a positive correlation between the CorrAdjust enrichment score and training set sample size (Spearman’s *r* = 0.85, *p* = 3.75 ∗ 10^−10^), with the lowest scores corresponding to datasets with < 50 samples. At the same time, the correlation is not linear and diminishes as sample size increases: the quadratic term *β*_2_ of the model score = *β*_0_ + *β*_1_ ∗ (sample size) + *β*_2_ ∗ (sample size) + *ϵ* is negative and statistically significant (*p* = 0.03), showing that an adequate sample is required for a successful CorrAdjust application.

### CorrAdjust outperforms other methods when correcting mRNA-mRNA correlations while eliminating fewer PCs

We next evaluated CorrAdjust and competing methods using mRNA-mRNA correlations and Canonical Pathways and GO as the reference collections (Figure 5). This time, we also analyzed 6,295 datasets from 11 healthy tissues from the GTEx dataset (bottom panels in Figure 5). We used *α* = 1% threshold for all these tests (see Supp. Note S1).

**Figure 5.**
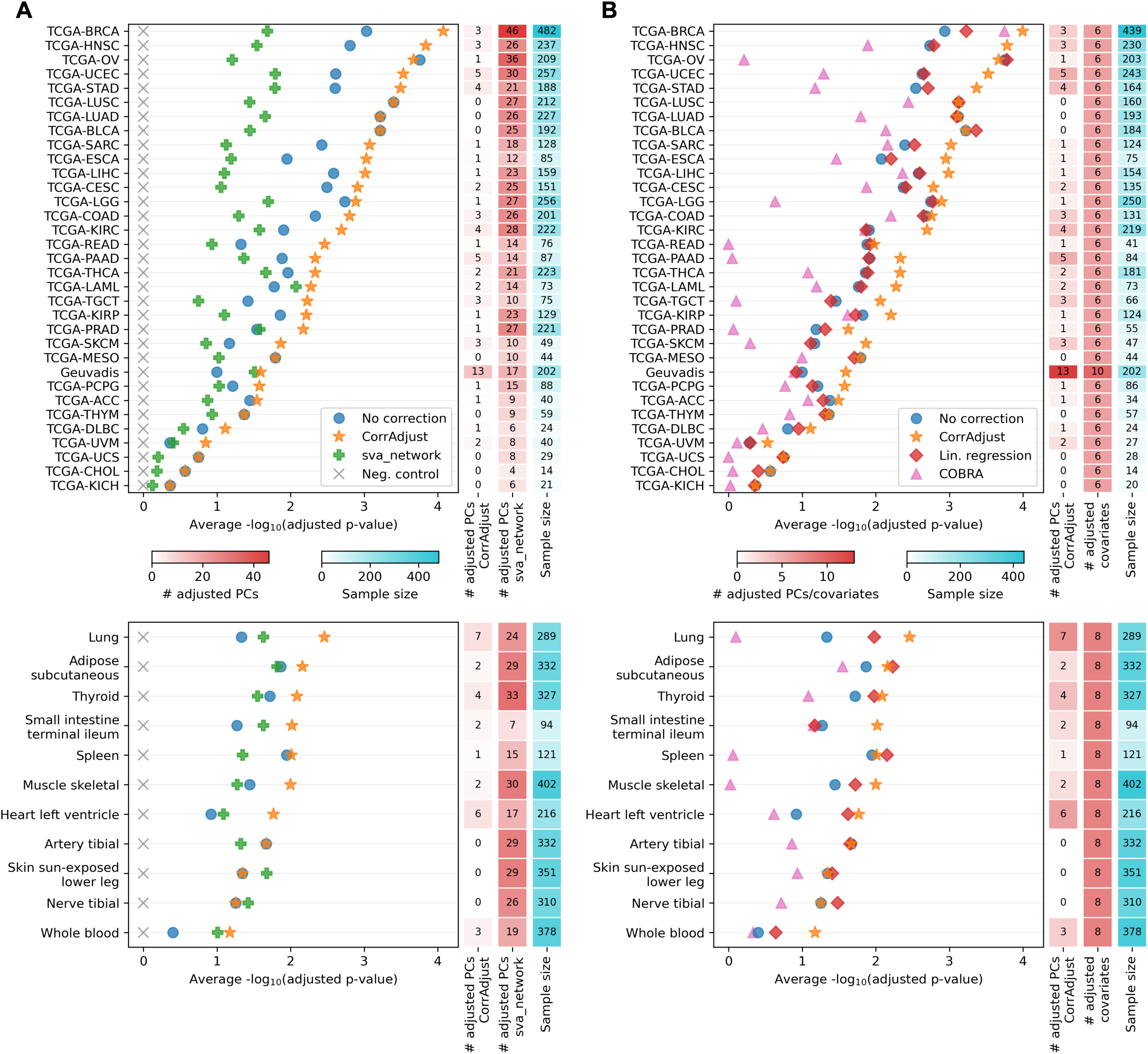
Benchmarking results of CorrAdjust and alternative methods using mRNA-mRNA correlations (test samples, all pairs). (A) Methods for hidden confounders correction. (B) Methods for known covariates correction. The top panels show TCGA and Geuvadis datasets, and the bottom ones show GTEx data. The X-axis shows the global enrichment score (average of Canonical Pathways and Gene Ontology). The columns to the right of the main plots show the number of PCs adjusted by CorrAdjust, the number of PCs adjusted by the sva_network approach, the number of adjusted known covariates, and the number of samples in the training set.

CorrAdjust again outperforms all competing methods when applied to the TCGA/Geuvadis (Figure 5; top panels) and GTEx (Figure 5; bottom panels) datasets. In almost all analyzed datasets, CorrAdjust marks only a few PCs as confounders (median = 1 PC). Interestingly, the differences between uncorrected and corrected values are not as high as in the miRNA-mRNA case. CorrAdjust enrichment score again positively and non-linearly correlates with the training set sample size (Spearman’s *r* = 0.35, *p* = 0.019; negative quadratic term *p* = 0.0067), with the lowest scores corresponding to datasets with < 50 samples.

Correcting for known confounders with linear regression has very little impact on enrichment scores for most datasets (Figure 5B). The sva_network and COBRA methods show significantly lower scores than uncorrected data in the vast majority of datasets, which is also the case with the training data (Supp. Figure S5).

## DISCUSSION

We developed CorrAdjust, a PCA-based method for correcting hidden confounders in transcriptomic correlation analyses. Confounder PC identification is guided by a novel enrichment-based metric designed to assess whether highly correlated gene pairs are enriched in user-specified reference pair sets (Figure 1). The metric is derived from individual gene scores, which can be visualized through volcano plots. These, along with other visualizations available in the CorrAdjust Python implementation, significantly enhance the interpretability of our approach (Figure 2, Figure 3). The corrected gene expression matrix produced by CorrAdjust can be used with any tools for downstream network analysis, such as WGCNA [4].

CorrAdjust clearly outperforms competitor methods for the correction of hidden and known confounders in the largest, to date, benchmark experiment. It includes over 25,000 human mRNA and small RNA sequencing datasets, spanning 32 cancer types from TCGA, 11 healthy tissues from GTEx, and LCLs from the Geuvadis consortium (Figure 4, Figure 5). Our collection covers all the GTEx tissues included in the previously reported mRNA-mRNA correlation benchmarks [11, 12, 14] (we are not aware of similar miRNA-mRNA studies). Consistently with the most recent report [12], correction for known covariates with linear regression has the least pronounced effect on correlations compared to the uncorrected data, while the sva_network method [14] performs worse than the uncorrected data for most datasets. The recently released COBRA tool [13] also shows scores mostly below the baseline.

At first sight, these data seem to differ from the results by Parsana et al., who reported significantly decreased FDR (equivalently, increased precision) with the GTEx data and Canonical Pathways collection upon sva_network correction [14]. The key difference between our analyses is that we evaluated raw and corrected correlations using the same number of <u>gene pairs</u> (*α*% pairs with the highest absolute correlations in each case). On the other hand, Parsana et al. evaluated gene pairs under the same WGCNA cut-height, essentially using the same <u>correlation cutoff</u>. In Supp. Note S2 we show that while doing so improves precision, it is incorrect because it ends up considering orders of magnitude fewer gene pairs since the sva_network correction strongly shrinks the correlations toward zero.

We observed the most profound impact of CorrAdjust on the enrichment scores with miRNA-mRNA correlations, where the reference pairs comprise miRNAs and their target mRNAs (Figure 4). Consistent with the existing literature [40–42], the baseline values for many of the datasets are very low. Geuvadis LCL data are the most extreme case, showing no miRNAs with a statistically significant overrepresentation of target mRNAs among the negative miRNA-mRNA correlations (Figure 3B). After applying CorrAdjust, 78% of the miRNAs in LCLs became statistically significant (Figure 3C), and the whole distribution of miRNA-mRNA correlations shifted to the biologically expected direction of negative correlations (Figure 3D-E). Although we used only miRNA targets to compute the scores, the corrected expression data can be used to derive correlations between other sncRNA types (e.g., tRFs, rRFs, or yRFs) and mRNAs to get insights about the functions of these much less studied molecules [38].

CorrAdjust differs from existing correlation correction methods by being reference-guided – it *selects* a subset of PCs in a supervised manner to distinguish between confounding and biological signals. In classical machine learning terms, CorrAdjust tackles a binary classification problem, assigning “shared” or “non-shared” labels to <u>gene pairs</u> based on a single feature: Pearson’s correlation. Note that <u>biological samples</u> are not *explicitly* referenced in this context, as their influence is inherently embedded within the correlations. To prevent overfitting and measure the generalizability of PC correction, we designed several specific methods. First, CorrAdjust automatically splits <u>gene pairs</u> (objects being classified) into 50% training and 50% validation sets, using the validation pairs for early stopping to determine the optimal number of PCs to adjust. We avoid the term “test set” for gene pairs, as the same samples are used for both PCA and correlations computation, thus compromising the independence of gene pairs. CorrAdjust also supports a <u>test set of samples</u> by re-using PC coefficients and identities derived from the training sample set. Finally, CorrAdjust includes an automated negative control experiment option that shuffles gene names while preserving expression data (and, thus, PCs and correlations) and reference collection structures. We observed no overfitting with all the used datasets in the benchmark experiment using all three techniques.

As with any correlation-based method, the main limitation of CorrAdjust is the requirement of a relatively large sample size; the benchmark runs showed reliable results when at least 50 samples were used. Another limitation of our analysis is the choice of reference collections. Here, we used Canonical Pathways and Gene Ontology to evaluate mRNA-mRNA correlations, despite neither of the resources positioning as a database of reference gene pairs for transcriptome-based correlation analysis. We avoided using the database of gene pairs GIANT [43], as one of the main contributors to its gene-gene connectivity score is “vanilla” Pearson’s correlation between expression levels computed over all Gene Expression Omnibus datasets, which might be heavily affected by common biases in transcriptomic analyses.

In conclusion, we believe confounder correction should be a standard practice for computing transcriptomic correlations. Such correction is especially vital when integrating different data modalities, such as the miRNA-mRNA case, as CorrAdjust increases the goodness of fit of correlations to experimentally validated data by orders of magnitude.

## Supporting information

CorrAdjust.Supplemental Material

## Author contributions

S.N., P.L., and I.R. designed CorrAdjust. S.N. performed theoretical analyses, implemented CorrAdjust, collected the data, and conducted experiments. S.N. and P.L. extensively tested CorrAdjust prior to its release. S.N. and I.R. wrote the manuscript with contributions from P.L. I.R. supervised the study. All authors approved the final version of the manuscript.

## Acknowledgments

This study was supported by Thomas Jefferson University Institutional Funds (I.R.), a National Foundation for Cancer Research (NFCR) grant (I.R.), a Melanoma Research Institute of Excellence (MRIE) at Thomas Jefferson University grant (I.R.), and NIH grants R21CA280575 (I.R.) and R01HG012784 (I.R.).

