## Supplementary material for "CorrAdjust unveils biologically relevant transcriptomic correlations by efficiently eliminating hidden confounders": CorrAdjust.Supplemental Material

### SUPPLEMENTAL FIGURE CAPTIONS

**Supp. Figure S1.** Evaluation of optimization trajectories derived from the training samples on the test samples (all gene pairs) for the GTEx whole blood (mRNA-mRNA) and Geuvadis LCL (miRNA-mRNA) datasets. Panels (A) and (B) correspond to the iterative residualization of PCs in the order shown in Figure 2A and Figure 3A, respectively.

**Supp. Figure S2.** Joint distributions of the identified confounder PCs and known covariates of the GTEx whole blood dataset (training samples). The data correspond to the correlations shown in Figure 2F.

**Supp. Figure S3.** Distribution of miRNA-mRNA correlations (test samples, all pairs) in the Geuvadis LCL datasets along the PC residualization trajectory (see Figure 3A). Solid lines correspond to the pairs of miRNAs and their experimentally validated targets from TarBase.

**Supp. Figure S4.** Benchmarking results of CorrAdjust and alternative methods using miRNA-mRNA correlations (training samples, training pairs). (A) Methods for hidden confounders correction. (B) Methods for known covariates correction. The X-axis shows the global enrichment score (average of TarBase and RNA22). The columns to the right of the main plots show the number of PCs adjusted by CorrAdjust, the number of PCs adjusted by the sva_network approach, the number of adjusted known covariates, and the number of samples in the training set. (C-E) The dependence of the global enrichment score (training samples) on the number of PCs removed by the sva_network approach (i.e., the k-th iteration corresponds to regressing out $PC_k$). Three representative datasets are shown (Geuvadis, TCGA-THCA, TCGA-PRAD).

**Supp. Figure S5.** Benchmarking results of CorrAdjust and alternative methods using mRNA-mRNA correlations (training samples, training pairs). (A) Methods for hidden confounders correction. (B) Methods for known covariates correction. The top panels show TCGA and Geuvadis datasets, and the bottom ones show GTEx data. The X-axis shows the global enrichment score (average of Canonical Pathways and Gene Ontology). The columns to the right of the main plots show the number of PCs adjusted by CorrAdjust, the number of PCs adjusted by the sva_network approach, the number of adjusted known covariates, and the number of samples in the training set.

**Supp. Figure S6.** Impact of parameter $\alpha$ on CorrAdjust performance on miRNA-mRNA correlations. (A) Histogram of the number of adjusted PCs across all TCGA cancer types and the Geuvadis collection. (B) Global enrichment scores before and after CorrAdjust correction (test samples, all pairs). Each marker represents one TCGA cancer type or the Geuvadis collection. P-values in the top-right corners were computed using paired sample Student's *t*-test.

**Supp. Figure S7.** Impact of parameter $\alpha$ on CorrAdjust performance on mRNA-mRNA correlations. (A) Histogram of the number of adjusted PCs across all TCGA cancer types, GTEx tissues, and the Geuvadis collection. (B) Global enrichment scores before and after CorrAdjust correction (test samples, all pairs). Each marker represents one TCGA cancer type, GTEx tissue, or the Geuvadis collection. P-values in the top-right corners were computed using paired sample Student's *t*-test.

**Supp. Figure S8.** Evaluation of the sva_network approach for correcting mRNA-mRNA correlations under different cutoffs. (A) Cumulative distribution functions of gene-gene correlations in the TCGA-BRCA data before and after sva_network correction (test samples, all pairs). Solid lines correspond to the pairs of mRNAs that are shared by at least one Canonical Pathway. (B) FDR scores for Canonical Pathways were evaluated at the four points shown in panel A (test samples, all pairs). Each marker represents one TCGA cancer type, GTEx tissue, or the Geuvadis collection. P-values were computed using paired sample Student's *t*-test. (C) The number of gene pairs corresponding to $\alpha = 1\%$ pairs for uncorrected data (X-axis) and the number of pairs corresponding to the same correlation thresholds after the sva_network correction (Y-axis). Each marker represents one TCGA cancer type, GTEx tissue, or the Geuvadis collection.

### SUPPLEMENTAL METHODS

#### Small RNA-seq read mapping

We used a modified version of our previously described pipeline to map the small RNA-seq reads [1]. Specifically, we profiled isomiRs and tRFs using IsoMiRmap [2] and MINTmap [3], respectively. We profiled rRFs, yRFs, and other repetitive fragments (rpFs) by mapping reads to rRNA, Y RNA, and RepeatMasker reference sequences, respectively, using an exhaustive, brute-force approach we described previously [1]. Next, we identified the unmapped reads with the Levenshtein distance ≤ 2 from at least one of the mapped small RNAs using the polyleven Python package (https://github.com/fujimotos/polyleven). We used the "license plates" labeling scheme [1-3] (https://cm.jefferson.edu/LicensePlates/) to uniformly label the mapped small RNAs using the prefixes "iso-," "tRF-," "rRF-," "yRF-," "rpF-," and "unk-" for isomiRs, tRFs, rRFs, yRFs, rpFs, and "Levenshtein distance ≤ 2" molecules, respectively.

Because we used reference collections of miRNA-mRNA interactions in this study, we let only the canonical isomiRs directly contribute to the CorrAdjust enrichment scores. However, we considered all small RNA types during the computation of PCs.

### SUPPLEMENTAL NOTES

#### Supplemental Note S1: The effect of parameter $\alpha$ on the performance of CorrAdjust

We ran CorrAdjust with six different values of $\alpha$ (0.1%, 0.5%, 1%, 5%, 10%, and 50%) across TCGA, GTEx, and Geuvadis datasets.

Supp. Figure S6 shows the effect of different $\alpha$ values on miRNA-mRNA correlations. Supp. Figure S6A shows the distributions of the number of PCs removed by CorrAdjust across TCGA and Geuvadis datasets. There are no significant differences between distributions for the values of $\alpha$ we considered. Supp. Figure S6B shows the distribution of global enrichment scores before and after the CorrAdjust application (each blue-point-and-orange-star pair stands for one TCGA cancer type or the Geuvadis dataset). At each value of $\alpha$, scores for corrected data are significantly higher than for the matched uncorrected data. At the same time, the scores rapidly grow until $\alpha = 5\%$, where they reach a plateau. We thus used $\alpha = 5\%$ to correct miRNA-mRNA data.

Supp. Figure S7 is a counterpart of Supp. Figure S6 for mRNA-mRNA correlations. Interestingly, at $\alpha \geq 5\%$, CorrAdjust removes no confounder PCs for most datasets, suggesting underfitting (Supp. Figure S7A). Concordantly, the differences in enrichment scores between uncorrected and CorrAdjust-corrected data are much more pronounced at $\alpha = 1\%$ and lower (Supp. Figure S7B). Thus, we used $\alpha = 1\%$ to correct mRNA-mRNA data.

#### Supplemental Note S2: Computing precision at a fixed *correlation cutoff* overestimates the performance of correction methods

We first illustrate the difference between our and sva_network [4] approaches for comparisons (fixed number of gene pairs vs. fixed correlation cutoff) using the TCGA-BRCA dataset. To be consistent with the data reported by Parsana et al. [4], we evaluate correlations using the FDR score (in the notation of Methods, $\mathrm{FDR} = 1 - \mathrm{P@}K$). Supp. Figure S8A shows the distribution of mRNA-mRNA correlations before correction and after removal of the first 46 PCs. The PC correction strongly shrinks the correlations toward zero. With our approach, we make a "horizontal" comparison between points 1 (corrected) and 2 (uncorrected) corresponding to the same value $\alpha = 1\%$. Consistently with Figure 5A, the FDR increases upon the sva_network correction (FDR = 0.83 after correction, FDR = 0.8 before correction). Following the approach of Parsana et al. [4], we make a "vertical" comparison of FDRs between points 2 (uncorrected) and 3 (corrected), corresponding to the same correlation cutoff of 0.53. Consistently with the original work, FDR decreases from 0.8 to 0.69 after the correction. However, point 3 corresponds to 58x fewer gene pairs than point 1 (7,833 vs. 452,629 pairs, respectively). Notably, when we evaluate uncorrected correlations of the top 7,833 gene pairs (point 4), FDR becomes even lower (0.57), and the sva_network approach again underperforms in the "horizontal" comparison (point 3 vs. point 4). Thus, in the case of TCGA-BRCA, the "vertical" comparison approach overestimates the performance of the PC correction by *incorrectly* trading off the number of gene pairs for the reduced FDR.

These results are not unique to the TCGA-BRCA example and generalize to the rest of the datasets. Supp. Figure S8B shows FDRs corresponding to the four points shown on Supp. Figure S8A: each marker stands for a TCGA cancer, a GTEx tissue, or the Geuvadis dataset. There is a modest *increase* in FDR upon the sva_network correction when the number of gene pairs is fixed (points 1 and 2). There is a substantial *decrease* in FDR upon the correction if the scores are computed at the same correlation cutoff (points 2 and 3). When the number of gene pairs is again equalized between uncorrected and corrected gene pairs, the improvement achieved by the sva_network procedure becomes much less pronounced (points 3 and 4). At the same time, the number of gene pairs that survive the same correlation cutoff (the vertical line containing points 2 and 3), on average, decreases by 234x after the correction (Supp. Figure S8C). Thus, the exceptional decrease in FDR score that Parsana et al. [4] observed is expected and results from the shrinkage of the correlation distributions toward zero coupled with leaving the correlation cutoff value unchanged, before and after the PC adjustment.

We also note that there is only a weak difference between the uncorrected data and the sva_network correction from the perspective of the FDR score (points 1 and 2 in Supp. Figure S8B). At the same time, Figure 5A shows a substantial decrease in global enrichment score upon correction for most datasets. This is explained by the fact that our enrichment score analyzes the neighborhood of each gene separately and thus can detect a decrease in the enrichment that affects many genes. On the other hand, FDR (or $\mathrm{P@}K$) analyzes all gene pairs together and fails to detect these differences adequately (see discussion of Simpson's paradox in Methods).

**Supplemental Note S3: Theoretical results**

**Lemma 1.** Let $X$ denote a gene expression matrix with $n$ rows (samples) and $d$ columns (genes). We assume that the columns of $X$ are centered. Let $u$ denote an $n \times 1$ vector and let $\tilde{X}$ denote a matrix derived by residualizing $u$ from each column of $X$. Finally, let $C$ and $\tilde{C}$ be the $d \times d$ covariance matrices of $X$ and $\tilde{X}$, respectively. Then, the first principal component (PC$_1$), normalized to unit length, is the solution to the following optimization problem:

$$\underset{\|u\|=1}{\operatorname{argmax}} \left\| C - \tilde{C} \right\|_F$$

Here, $\|\cdot\|_F$ denotes the Frobenius norm of a matrix.

**Proof.** Let $\beta$ be a $1 \times d$ vector of coefficients from regressing the columns of $X$ on $u$, i.e., $\tilde{X} = X - u\beta$. The columns of the residual matrix $\tilde{X}$ are orthogonal to $u$, meaning $\tilde{X}^T u = 0$. Consequently, we have $X^T u = \beta^T u^T u = \beta^T$. By the definition of the covariance matrix, maximizing $\left\| C - \tilde{C} \right\|_F$ is equivalent to maximizing $\left\| X^T X - \tilde{X}^T \tilde{X} \right\|_F$. Using the above formulas,

$$X^T X - \tilde{X}^T \tilde{X} = X^T X - \tilde{X}^T (X - u\beta) = X^T X - \tilde{X}^T X = \beta^T u^T X = X^T u u^T X$$

Both $X^T u$ and $u^T X$ are $d$-dimensional vectors, allowing us to reduce the Frobenius norm to the standard Euclidean norm:

$$\|X^T u u^T X\|_F = \|X^T u\|^2 = u^T X X^T u$$

Maximizing this expression with respect to the unit vector $u$ is a classical formulation for variance maximization, which corresponds to finding the PC$_1$ *coefficients* for the matrix $X^T$ (not $X$ itself). From the singular value decomposition (SVD) of $X$ and $X^T$, we see that $u$ represents both the first right singular value of $X^T$ (the PC$_1$ coefficients of $X^T$) and the first left singular value of $X$ (the PC$_1$ of $X$, scaled to unit length):

$$X = U \Sigma V^T, X^T = V \Sigma U^T$$

Plugging the SVD of $X$ and the optimal $u$ into the maximized function shows that $\left\| X^T X - \tilde{X}^T \tilde{X} \right\|_F$ equals the square of the first singular value, $\Sigma_1^2$.

Note that Lemma 3 can also be applied to the residuals $\tilde{X}$ after removing the effect of PC$_1$, showing that regressing out PC$_2$ from $\tilde{X}$ maximizes the change in the covariance matrix $\tilde{C}$. This Lemma can be applied iteratively to find subsequent principal components.

**Lemma 2.** Let $N$ denote the total number of samples and $n$ denote the total number of positive samples. Suppose that a classification algorithm predicts an $\alpha$-fraction of all samples as positives. Then, the following inequality holds:

$$\text{sensitivity} \leq \alpha \frac{N}{n}$$

**Proof.** Let $k$ be the number of true positives among the $\alpha N$ samples predicted as positive by the classifier. Clearly, $k$ cannot exceed $\alpha * N$, since this is the total number of samples the classifier predicted as positives. Thus, we have:

$$\text{sensitivity} = \frac{k}{n} \leq \alpha \frac{N}{n}$$

**Lemma 3.** Let $N$ denote the total number of samples, $n$ denote the total number of positive samples, $K$ denote the number of predicted positives, $k$ denote the number of true positives among the $K$ predictions. Then, the balanced precision (BP) [5] could be expressed as:

$$\text{BP} = \frac{\text{enr} * (1 - \pi)}{\text{enr} * (1 - 2\pi) + 1},$$

where $\text{enr} = \frac{k * N}{K * n}$ and $\pi = n/N$.

**Proof.** By the original definition (Equation 1 in [5]), the balanced precision is

$$\text{BP} = \frac{\frac{k}{K} * (1 - \pi)}{\frac{k}{K} * (1 - \pi) + \left(1 - \frac{k}{K}\right) * \pi}$$

Manipulating the expression, we get,

$$\text{BP} = \frac{1 - \pi}{1 - \pi + \left(\frac{K}{k} - 1\right) * \pi} = \frac{1 - \pi}{1 - \pi + \left(\frac{1}{\text{enr} * \pi} - 1\right) \pi} = \frac{1 - \pi}{1 - 2\pi + 1/\text{enr}} = \frac{\text{enr} * (1 - \pi)}{\text{enr} * (1 - 2\pi) + 1}$$
