## Supplementary figures and images for "CorrAdjust unveils biologically relevant transcriptomic correlations by efficiently eliminating hidden confounders"

### Supp. Figure S1

Supp. Figure S1

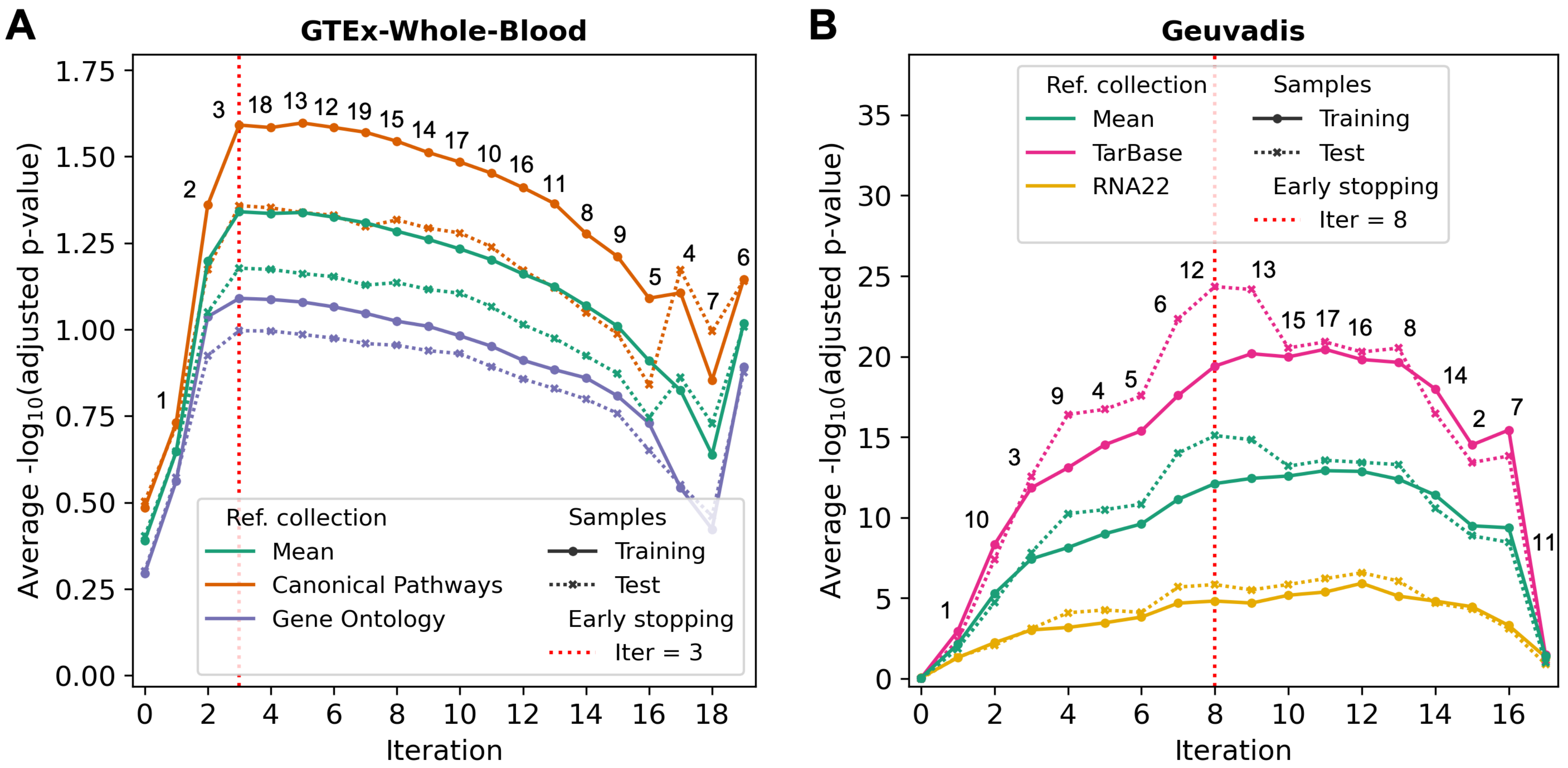

### Supp. Figure S2

#### Supp. Figure S2

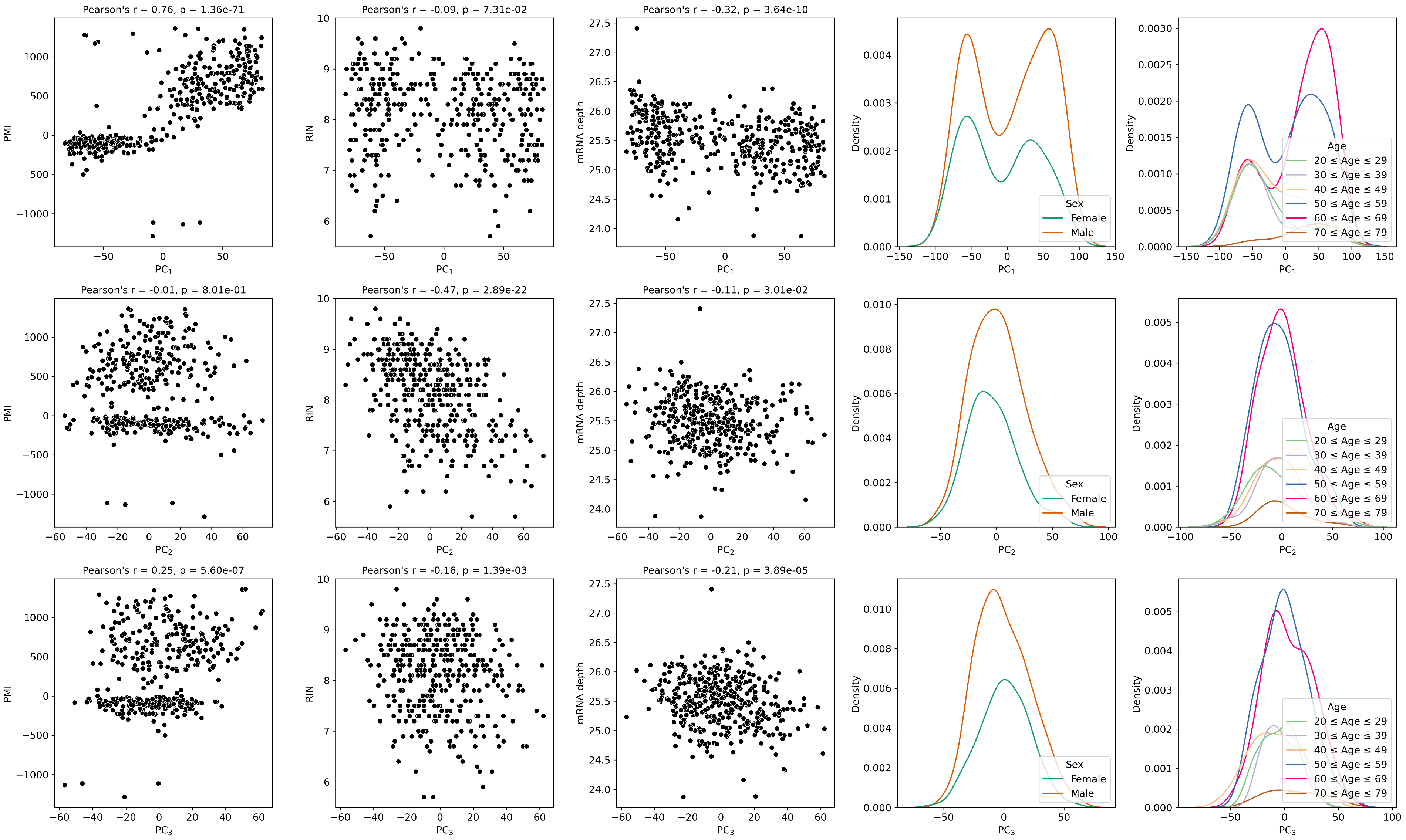

### Supp. Figure S3

Supp. Figure S3

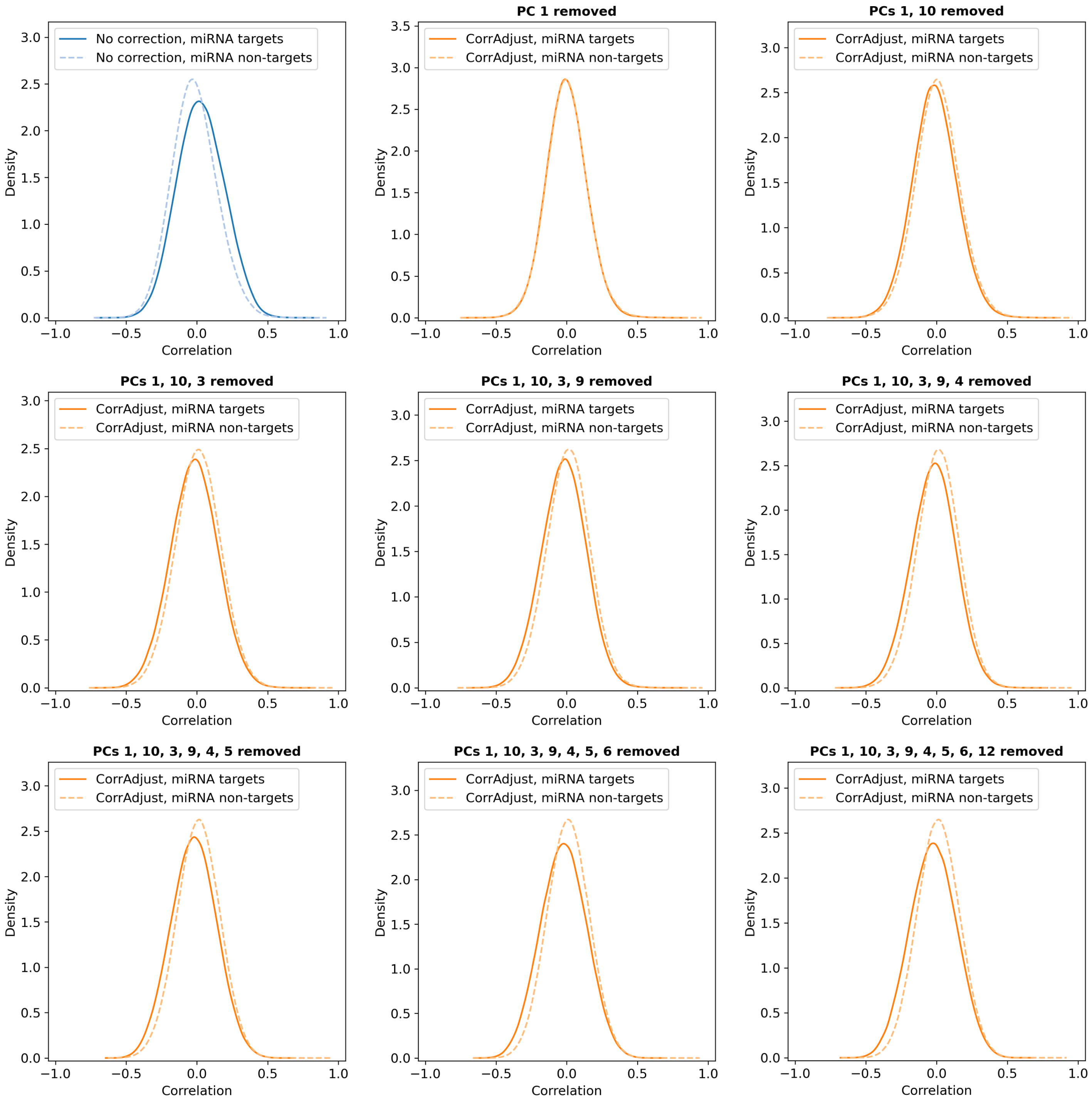

### Supp. Figure S4

Supp. Figure S4

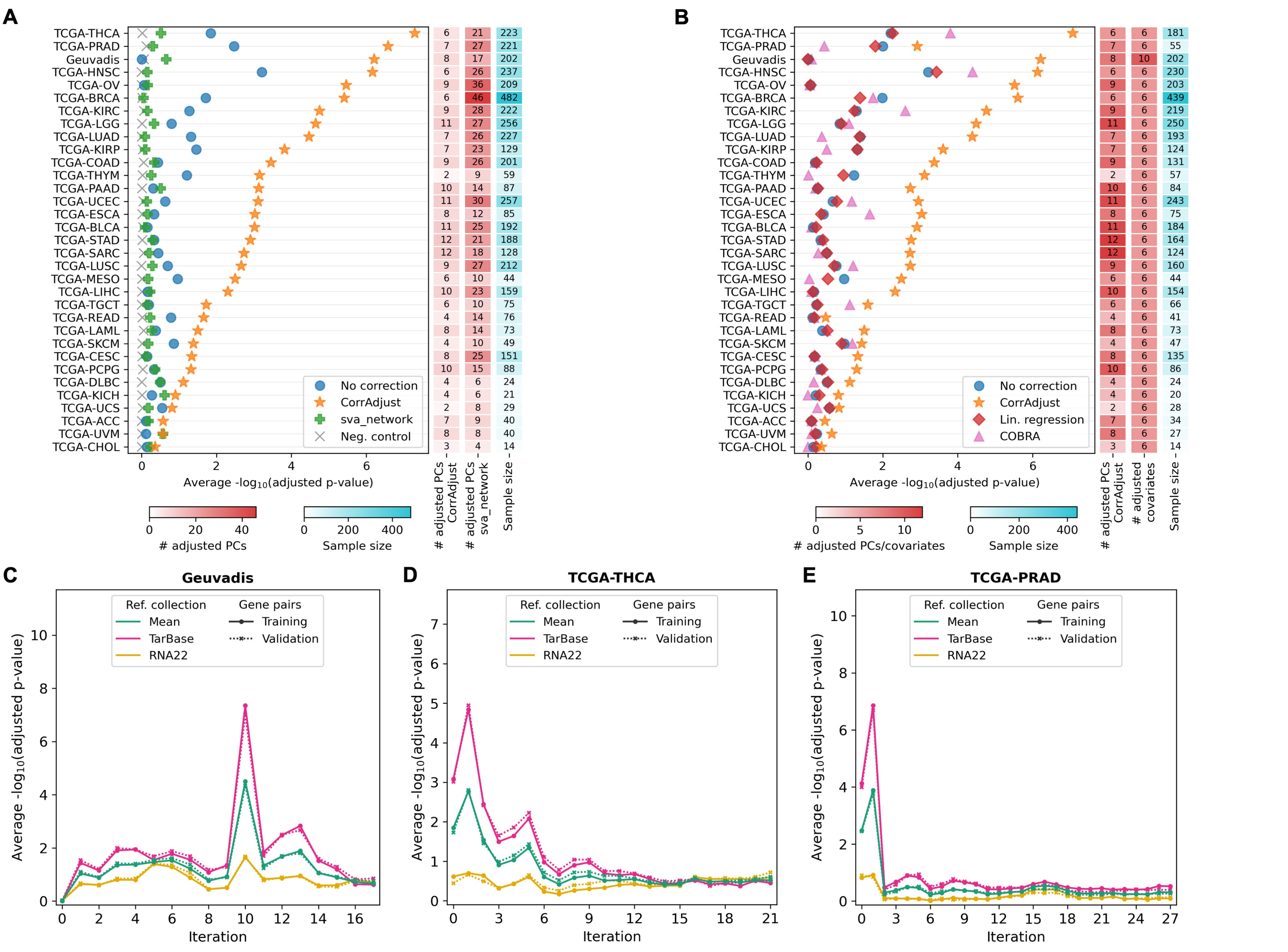

### Supp. Figure S5

Supp. Figure S5

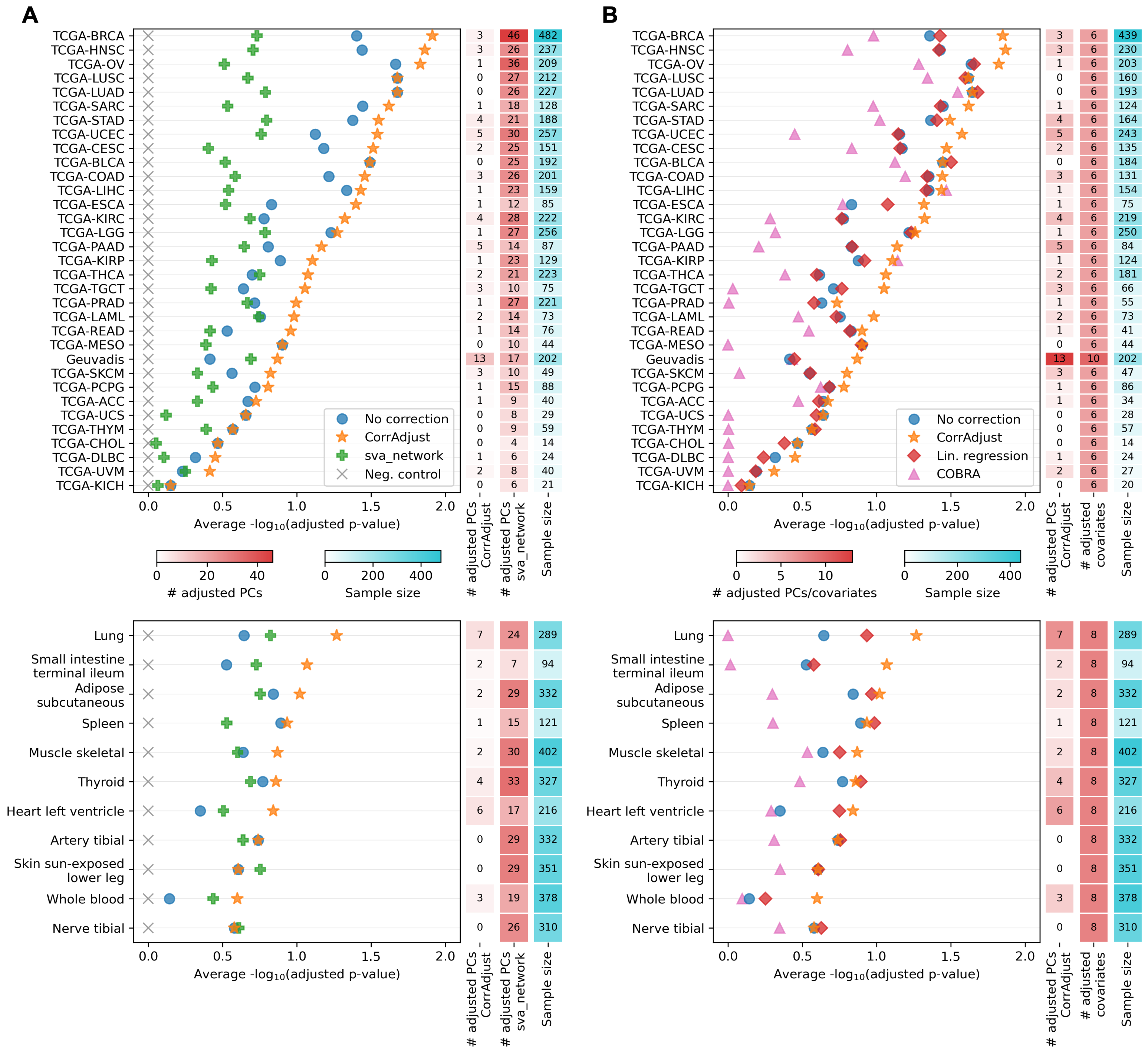

### Supp. Figure S6

Supp. Figure S6

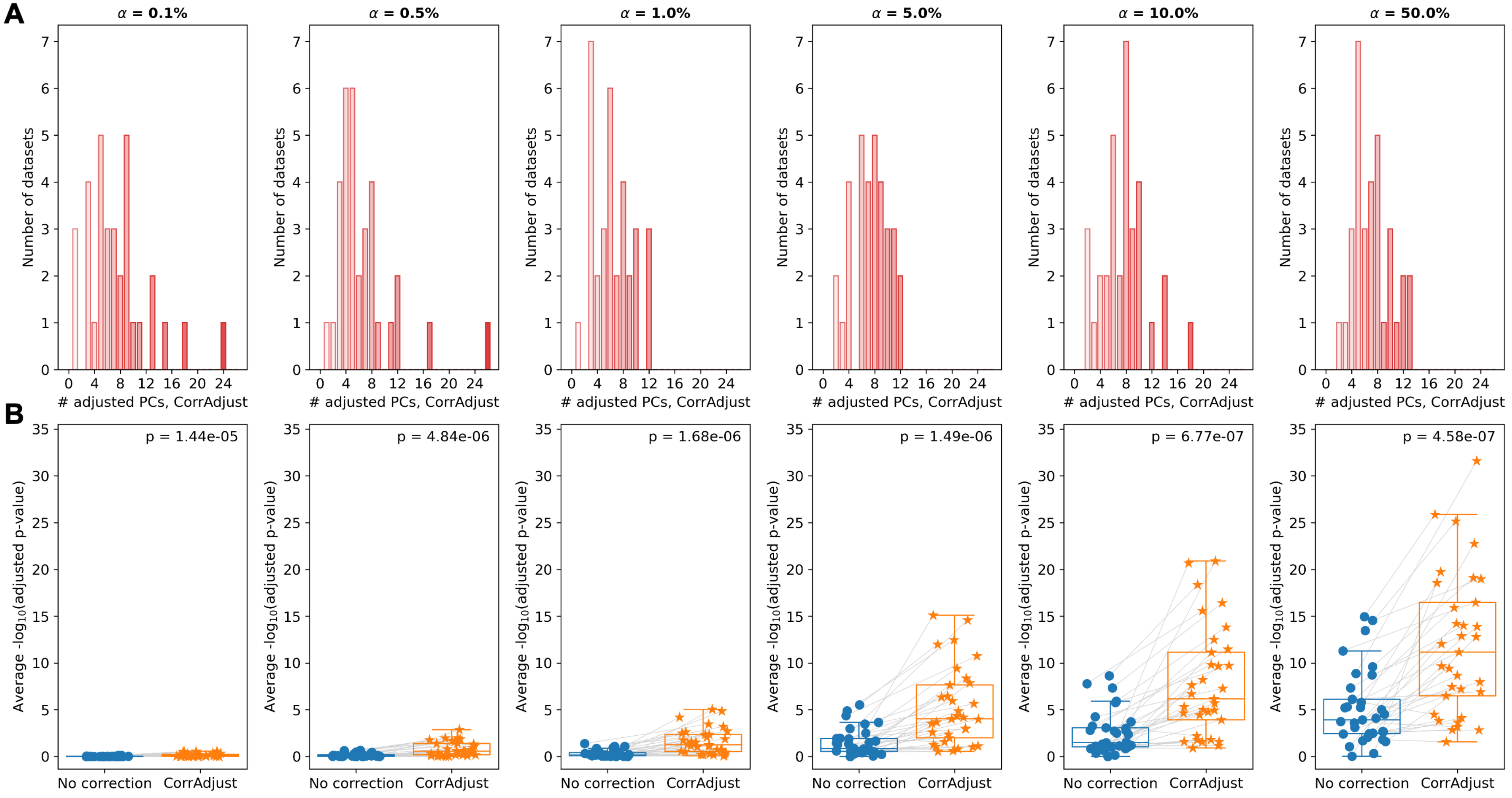

### Supp. Figure S7

Supp. Figure S7

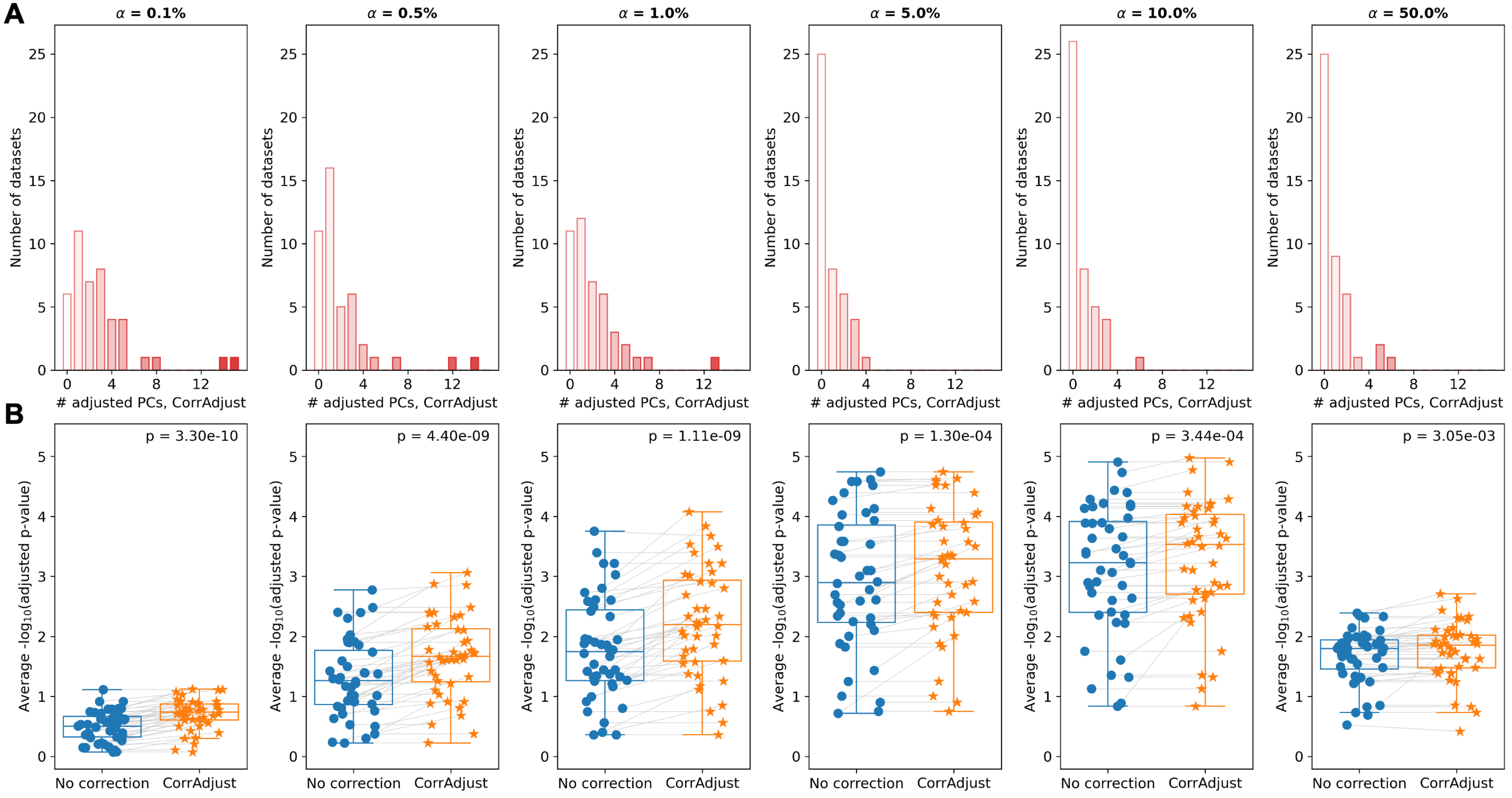

### Supp. Figure S8

Supp. Figure S8

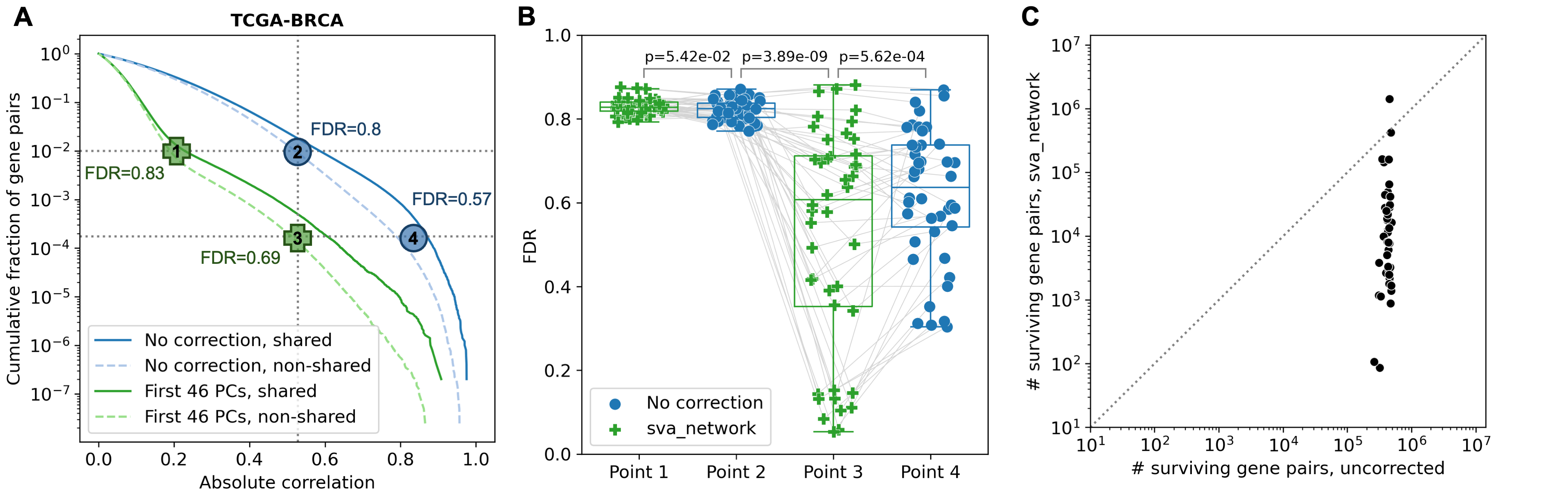
